# CNCDatabase: a database of non-coding cancer drivers

**DOI:** 10.1101/2020.04.29.069047

**Authors:** Eric Minwei Liu, Alexander Martinez-Fundichely, Rajesh Bollapragada, Maurice Spiewack, Ekta Khurana

## Abstract

Most mutations in cancer genomes occur in the non-coding regions with unknown impact to tumor development. Although the increase in number of cancer whole-genome sequences has revealed numerous putative non-coding cancer drivers, their information is dispersed across multiple studies and thus it is difficult to bridge the understanding of non-coding alterations, the genes they impact and the supporting evidence for their role in tumorigenesis across multiple cancer types. To address this gap, we have developed CNCDatabase, Cornell Non-Coding Cancer driver Database (https://cncdatabase.med.cornell.edu/) that contains detailed information about predicted non-coding drivers at gene promoters, 5’ and 3’ UTRs (untranslated regions), enhancers, CTCF insulators and non-coding RNAs. CNCDatabase documents 1,111 protein-coding genes and 90 non-coding RNAs with reported drivers in their non-coding regions from 32 cancer types by computational predictions of positive selection in whole-genome sequences; differential gene expression in samples with and without mutations; or another set of experimental validations including luciferase reporter assays and genome editing. The database can be easily modified and scaled as lists of non-coding drivers are revised in the community with larger whole-genome sequencing studies, CRISPR screens and further experimental validations. Overall, CNCDatabase provides a helpful resource for researchers to explore the pathological role of non-coding alterations and their associations with gene expression in human cancers.

## INTRODUCTION

Mutations in the cancer genome can be divided into drivers and passengers. Driver mutations are the ones that confer selective advantage for the cancer cells to grow. Multiple databases collecting protein-coding driver mutations in cancers, such as COSMIC, Intogen, OncoKB, and CIViC, have helped further follow-up investigations and enabled the utility of the driver catalogue in numerous basic and translational research studies (1–4). Recent studies have shown that besides mutations in protein-coding regions, mutations in non-coding regions, such as promoters, enhancers, insulators, and non-coding RNAs can also act as cancer drivers (5–12). Although mutations at the *TERT* promoter are the most prominent example of non-coding drivers, evidences supporting the functional role of other non-coding mutations as cancer drivers are dispersed in several independent publications. Different computational approaches and experimental methods have used different signals to identify non-coding cancer drivers and it is hard to assess their consensus in the absence of a unified database. The lack of a database dedicated to non-coding cancer drivers hinders a comprehensive evaluation of the number of drivers identified for their further downstream computational analysis, functional characterization, and their utility for translational research. Here, we built CNCDatabase (**C**ornell **N**on-coding **C**ancer driver database), a manually curated database that contains detailed information of non-coding cancer drivers from published studies. Currently, the CNCDatabase contains 1201 genes with significant alterations in the non-coding regions in 31 cancer types from 25 published articles.

## MATERIALS AND METHODS

### Data model

The CNCDatabase has been designed as a relational database to store the collected non-coding cancer drivers from multiple sources. The detailed Entity-Relationship (ER) diagram and description of all tables are provided on the “Download” page of the CNCDatabase website. The database schema follows a snowflake structure where the multidimensional data is connected to the centralized fact table. The design follows the database normalization rules for keeping the data integrity of multiple related entities such as: non-coding driver evidence, functional element and gene associations, cancer type and reported study (Figure 1A and supplementary figure 1). The data structure employed in the CNCDatabase allows it to be extended to accommodate new types of data without major changes in the existing model. As a result, the database is highly scalable which is a key feature for early-stage projects in data integration.

**Figure 1.**
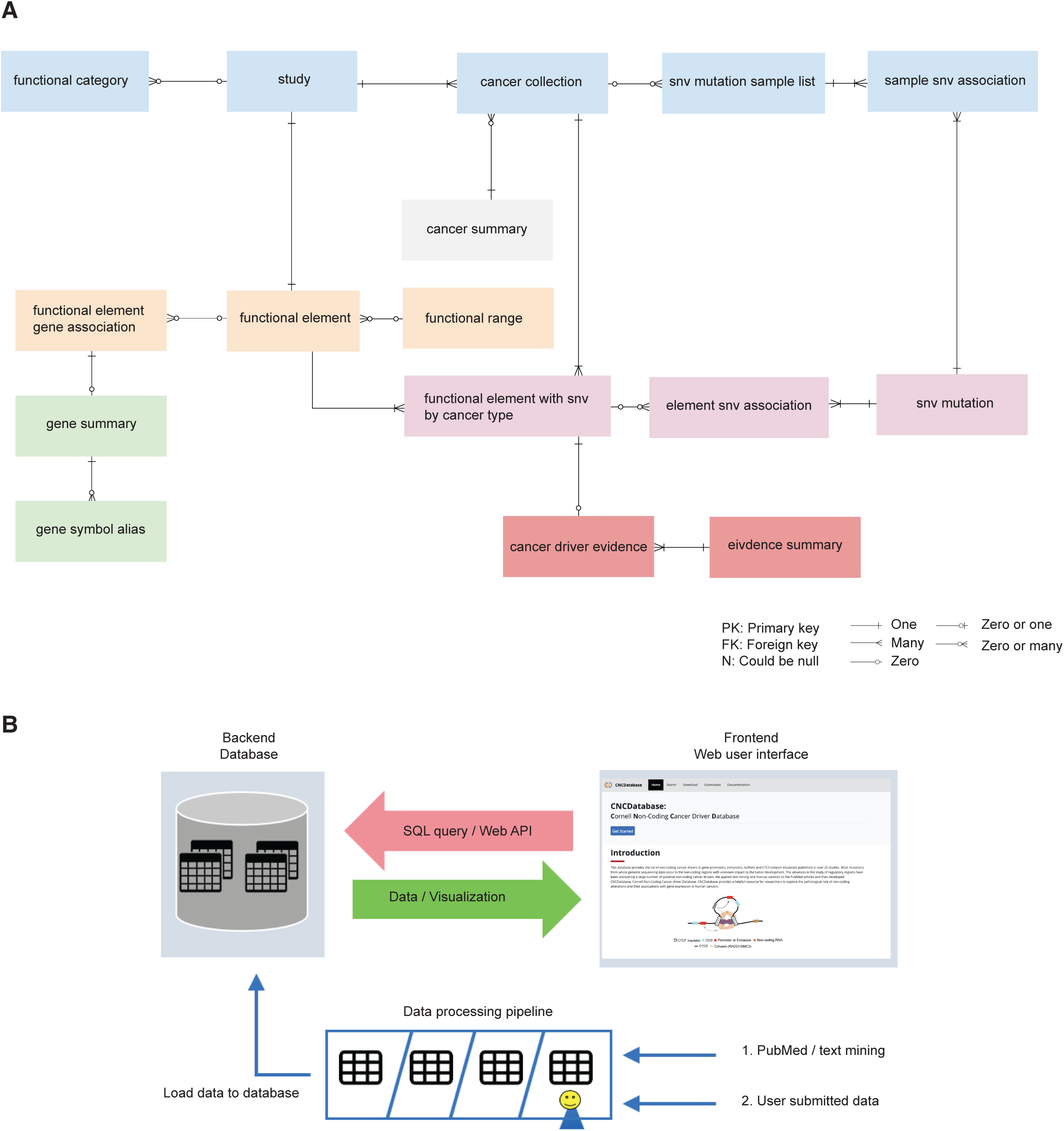
Data model and architecture of CNCDatabase. **(A)** Simplified database entity-relationship diagram (ERD). **(B)** The schematic data flow in the CNCDatabase between web interface in the frontend and PostgreSQL database in the backend. Manually curated cancer driver lists from PubMed or from users can be loaded into the database.

### Data collection and processing

We comprehensively gathered all studies related to cancer non-coding drivers currently available in the PubMed database up to 10^th^ February 2020. We text mined within the title or abstract of the articles for the existence of combinations of key words such as noncoding, driver, cancer, and search for their alternative terms. example: noncoding[Title/Abstract]) OR non-coding[Title/Abstract]) AND driver[Title/Abstract]). After manual review of the returned abstracts from PubMed search, we extracted the non-coding driver evidences in the text and supplementary files of the 25 selected articles. We focused on the publications reporting non-coding alterations in the promoters, 5’ UTRs, 3’ UTRs, enhancers, splice sites, non-coding RNAs and CTCF-cohesin insulators.

Because the CNCDatabase aims to catalogue the comprehensive list of human non-coding cancer drivers, we include the ones with at least one type of evidence: computational prediction, differential gene expression association from RNA-seq and other experimental validation. The evidence term “computational prediction” means the non-coding regions exhibit statistically significant signals of positive selection from whole genome sequencing data. The term “differential gene expression association from RNA-seq” means the mutations in the non-coding region are associated with differential gene expression between wild type and mutated samples from RNA-seq data. Finally, “other experimental validation” means the mutations in the non-coding region have been validated for molecular or cancer-related phenotype by either luciferase assay, CRIPSR-Cas9 or some other experimental assays.

### Architecture of CNCDatabase

CNCDatabase consists of a relational database server using PostgreSQL (version 9.6.6). It provides an application program interface (API) to access all stored data. The backend server is complemented with a frontend web-based user interface (UI) (Figure 1B). We uses Node.js (version 10.15.3) and Express.js framework (version 4.16.4) to build the backend server. The backend server also provides representational state transfer (REST) API so that the data can be accessed programmatically by other external web services. We uses React.js (version 16.8.5) and Bootstrap4 (version 4.0.0) as the frontend web development framework for responsive user interface, which means the website is suitable for both desktop and mobile data viewing. The chart visualizations are implemented by using plotly.js (version 1.46.1) package.

The CNCDatabase is freely available (https://cncdatabase.med.cornell.edu/). The content in the database is also available to download. We provide the code at GitHub (https://github.com/kuranalab/CNCDatabase) for users to make use of all services locally.

## DATABASE FEATURES AND USE

### Summary of database content

A total of 1676 entries in CNCDatabase correspond to 90 non-coding RNAs and non-coding regions of 1111 protein-coding genes that are associated with significant alterations in the non-coding regions from computational predictions, 18 genes associated with differential gene expressions from RNA-seq and 21 genes with other experimental validations in 32 cancer types (Figure 2). Out of the 1201 genes associated with non-coding drivers from computational predictions, 355 genes associated with predicted non-coding drivers are from individual cancer type analysis and 684 genes associated with predicted non-coding drivers are from pan-cancer analysis only where samples from multiple cancer types are pooled together for statistical power. The number of genes for individual cancer types varies from 270 in melanoma to 1 in RHAB (Figure 2A). The publication from Weinhold et al. contributes the largest number of non-coding candidates (453 genes) from computational predictions (13).

**Figure 2.**
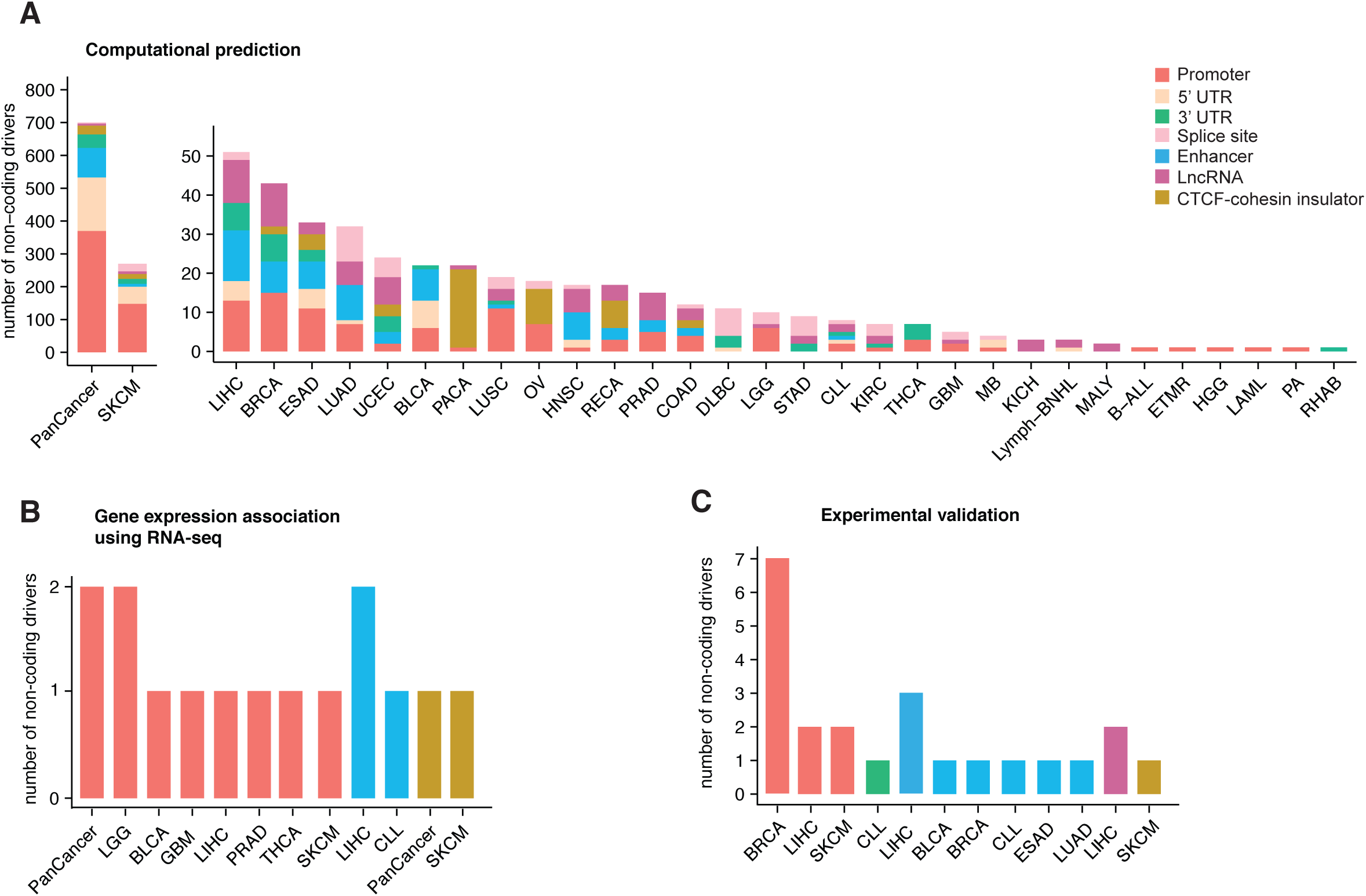
Summary of number of non-coding drivers. **(A)** Number of non-coding drivers in each cancer type by computational prediction. Cancer types include pan-cancer (PanCancer), skin cutaneous melanoma (SKCM), liver hepatocellular carcinoma (LIHC), breast invasive carcinoma (BRCA), esophageal carcinoma (ESAD), lung adenocarcinoma (LUAD), uterine corpus endometrial carcinoma (UCEC), bladder urothelial carcinoma (BLCA), pancreatic ductal adenocarcinoma (PACA), lung squamous cell carcinoma (LUSC), ovarian serous cystadenocarcinoma (OV), head and neck squamous cell carcinoma (HNSC), kidney renal papillary cell carcinoma (RECA), prostate adenocarcinoma (PRAD), colon adenocarcinoma (COAD), lymphoid neoplasm diffuse large B-cell lymphoma (DLBC), low grade glioma (LGG), stomach adenocarcinoma (STAD), chronic lymphoctytic leukemia (CLL), kidney renal clear cell carcinoma (KIRC), (THCA), glioblastoma multiforme (GBM), medullablastoma (MB), kidney chromophobe (KICH), (Lymph-BNHL), malignant lymphoma (MALY), B-cell acute lymphoblastic leukemia (B-ALL), embryonal tumor with multilayered rosettes (ETMR), high-grade glioma (HGG), (LAML), pilocytic astrocytoma (PA), rhabdoid tumor (RHAB). **(B)** Number of non-coding drivers in each cancer type that show differential gene expression in samples with mutations vs. without using RNA-seq data. **(C)** Number of non-coding drivers in each cancer type with support from other functional validation, such as CRISPR/Cas9 or luciferase reporter assay.

### Web user interface

CNCDatabase provides intuitive web interface that facilitates browsing and searching through four main sections including “Home”, “Search”, “Download”, “Submission” and “Documentation” (Figure 3). The landing page (“Home”) provides abstract graphics of the current available information. From there a simple button (“Get started”) immediately allows to launch user’s custom query. All data in the CNCDatabase can be downloaded from the “Download” section as text format files or database contents for further downstream analysis.

**Figure 3.**
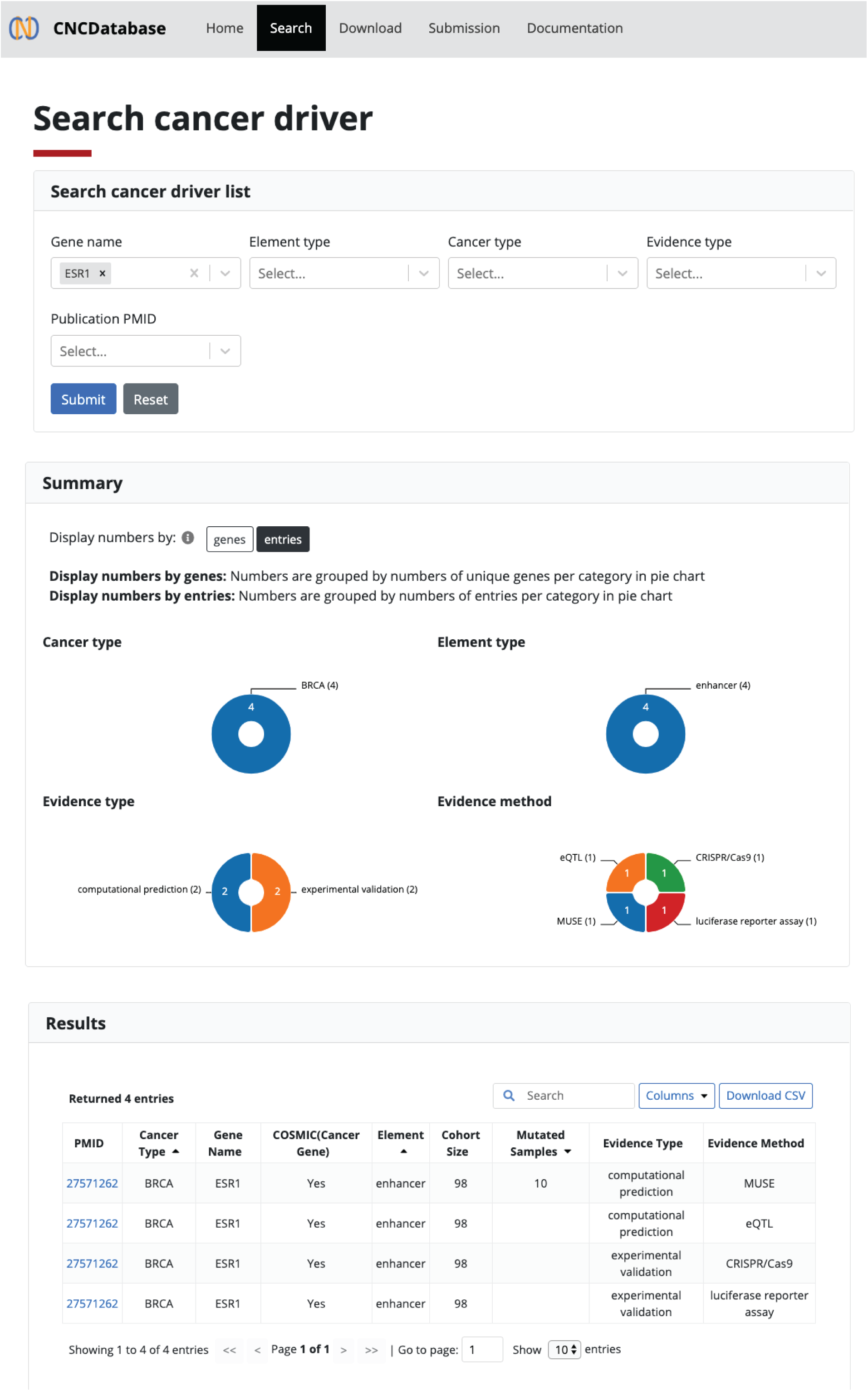
Web use interface and supported functionality in the CNCDatabase. User can use the combination of gene name, element type, cancer type, evidence type or publication id (PMID) to query the non-coding cancer driver list from the backend database. The returned result shows graphical summary and list in the table format.

#### Searching for non-coding cancer drivers

In the “Search” section, users can apply the fuzzy query to retrieve the non-coding driver entries. The query fields also support the auto-complete function so that users can quickly pick a valid gene name and cancer type. The database can be searched using multiple query types, including gene name, element type (e.g. promoter or enhancer), cancer type, evidence type and publication PMID. If users do not select a specific cancer type, the system will return results in all cancer types by default. After clicking “Submit” button, the query results are displayed in a report organized into several components. The “Summary” section provides pie chart representations to display the numbers in each category including cancer type, element type, evidence type and evidence method. The “Results” section displays the retrieved entries in tabular format including publication id (PMID), cancer type, gene name, Cancer Gene Census (CGC) from the catalogue of somatic mutations in cancer (COSMIC), non-coding element type, cohort size, mutated sample size, evidence type, and evidence method. Users can also further refine the search results by entering the targeted value in the search field of the returned result table (Figure 3). The search results can also be downloaded in the CSV format.

#### Data submission and curation

Through the “Submission” page of the web interface, users can submit new non-coding cancer driver evidences to the CNCDatabase. A valid data submission will need the user to prepare the content following the pre-defined text format that describes the publication id, cancer type, gene name, non-coding element type, cohort size, mutated sample size, evidence type, and evidence method. Then, users will receive separate email notifications to track the progress and correctness of their data submission. Thus, the CNCDatabase can serve as a central hub of non-coding cancer drivers’ exchange for the cancer research community regardless of users’ bioinformatics expertise level.

### Overview of data in CNCDatabase

One of the many uses of CNCDatabase is that it will help researchers to prioritize the non-coding candidates for functional validation follow-up and to look up which non-coding mutations have already had functional validation evidence.

Analysis of data in CNCDatabase reveals that the promoters of *TERT, WDR74, PLEKHS1* and *CCDC107* have support as non-coding drivers by computational predictions from more than four publications (Figure 4A). *TERT, WDR74* and *PLEKHS1* promoter mutations also have supporting evidence by gene expression in RNA-seq or by other experimental assays. It will be interesting to interrogate the function of *CCDC107* promoter mutations in breast, lung and rectal cancers in future studies (Figure 4B). In the 3’UTR regions, only *NOTCH1* in CLL has functional assay evidence (Figure 5A). *API5, DRD5, FAM230A* and *PCMTD1* could be good candidates for follow-up functional validations at 3’UTR regions. While many studies have identified candidate drivers at enhancers, *TP53TG1* is the only gene that has support from multiple publications. In the enhancer regions, the validation results are dispersed in separate publications (Figure 5B). For lncRNAs, *MALAT1* and *NEAT1* are the genes with support both from computational predictions and from functional assays (Figure 5C). Although mutations in the 5’UTRs and splice sites do not have any support as cancer drivers from functional assays in any published study yet, there are multiple genes (*WDR74, C16orf59, MED31, MTG2, PTDSS1, TBC1D12*, and *UMPS*) with support from computational predictions from multiple publications (Figure 5D). The splice site mutations of *TP53* and *STK11* are the most promising candidates to conduct follow-up validations (Figure 5E).

**Figure 4.**
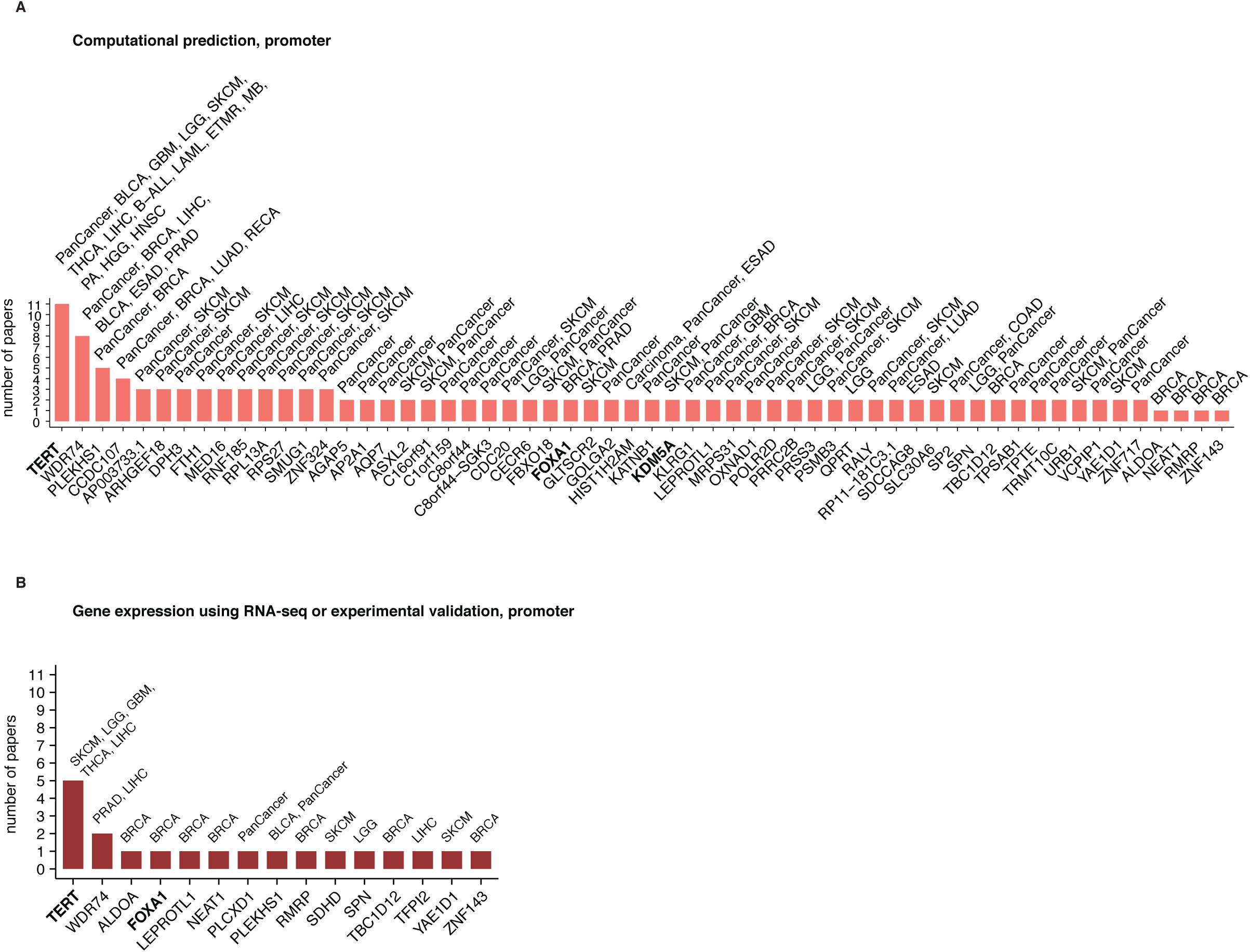
Non-coding cancer driver candidates at promoter regions. **(A)** computational predictions **(B)** gene expression association and other experimental validations. In the results from computational predictions, for cancer driver candidates reported in only one publication, we only show candidates with support from experimental validation or those associated with cancer genes in COSMIC census list. The cancer genes are highlighted in bold.

**Figure 5.**
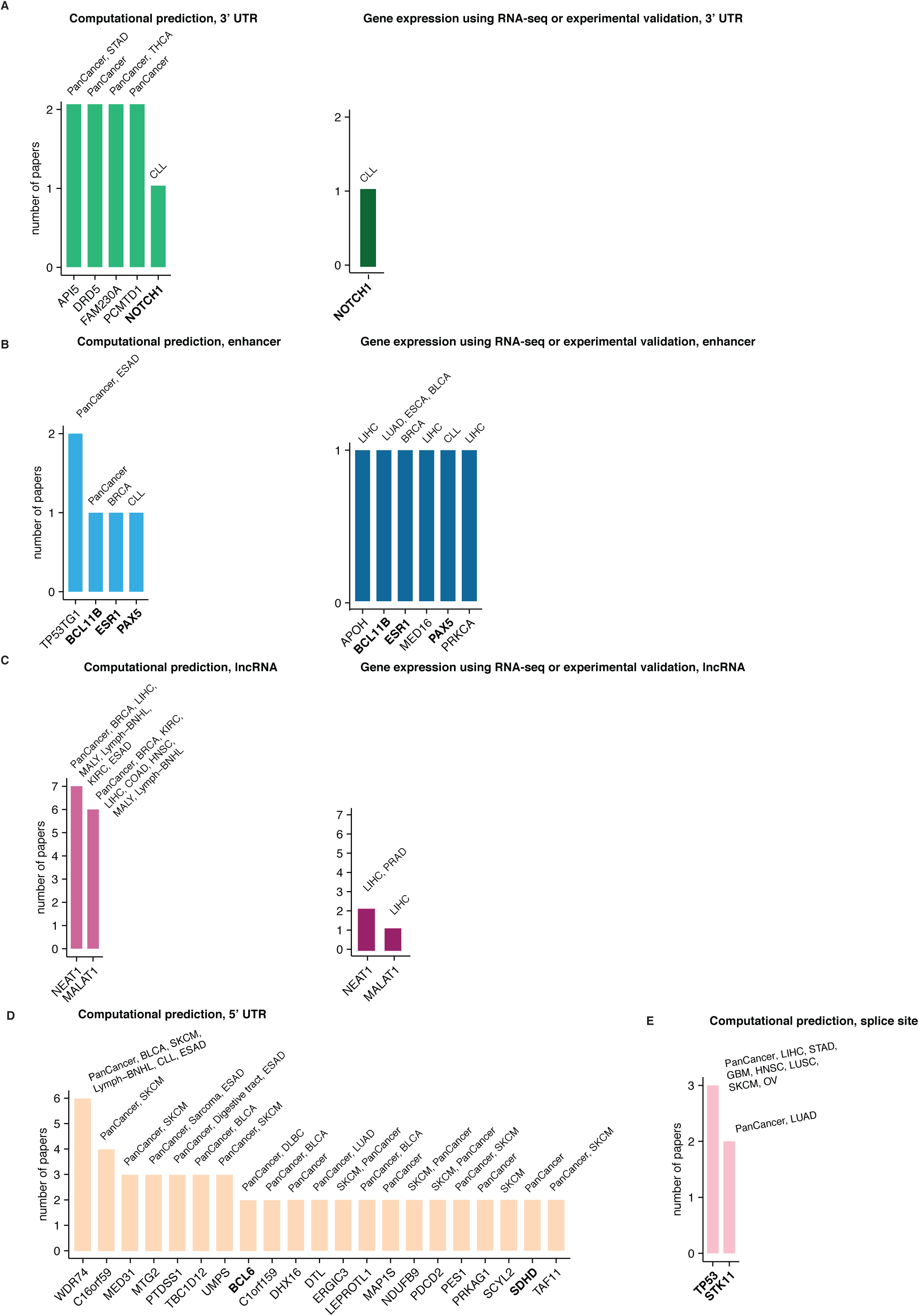
Non-coding cancer driver candidates from computational predictions and candidates with functional validations. **(A)** 3’ UTR, **(B)** enhancer, **(C)** lncRNA, **(D)** 5’UTR and **(E)** splice site. In the results from computational predictions, for cancer driver candidates reported in only one publication, we only show candidates with support from experimental validation or those associated with cancer genes in COSMIC census list. The cancer genes are highlighted in bold.

## DISCUSSION AND FUTURE PERSPECTIVES

We report the CNCDatabase that integrates the functional evidence reported for non-coding cancer drivers in many independent publications. To create this comprehensive catalogue, we used combinations of keywords to select relevant articles from PubMed and manually extracted the evidences hidden in the supplementary data of each article. At the time of writing this publication, we document 1300 non-coding cancer drivers with support from either computational prediction, gene expression association or other experimental validation. Our database aims to advance the understanding of non-coding alterations in cancer for both basic and translational scientists and users with all levels of bioinformatics skills. Novice users can use interactive queries to browse the evidences supporting non-coding cancer drivers and export the search results for further custom analysis. Users with advanced programming skills can use the RESTful APIs of the CNCDatabase in conjunction with their current analysis pipelines.

We will update our data collection periodically to incorporate the rapidly accumulating large-scale genomics studies of non-coding cancer drivers. With the advances in CRISPR screening technology, we expect more functionally validated non-coding cancer drivers will be reported in the future. In fact, CNCDatabase can help scientists pick the relevant lists of non-coding alterations for CRISPR validations whose results can be then added to the database to augment the functional evidence supporting or rejecting those drivers. In conclusion, CNCDatabase will serve as a valuable resource to complement the studies of oncogenic mechanisms currently centered on protein-coding mutations by the majority of cancer community.

## ACKNOWLEDGEMENTS

We thank Douglas Duckworth from Weill Cornell Supercomputing Unit for help in the website deployment. We also thank Ann Palladino, Sandra Cohen, Tawny Cuykendall, Erica Duo Xu and Andre Forbes at the Khurana lab for valuable suggestions.

## FUNDING

E.K. thanks the National Institutes of Health for support.

## FIGURE LEGENDS

**Supplementary figure 1.**
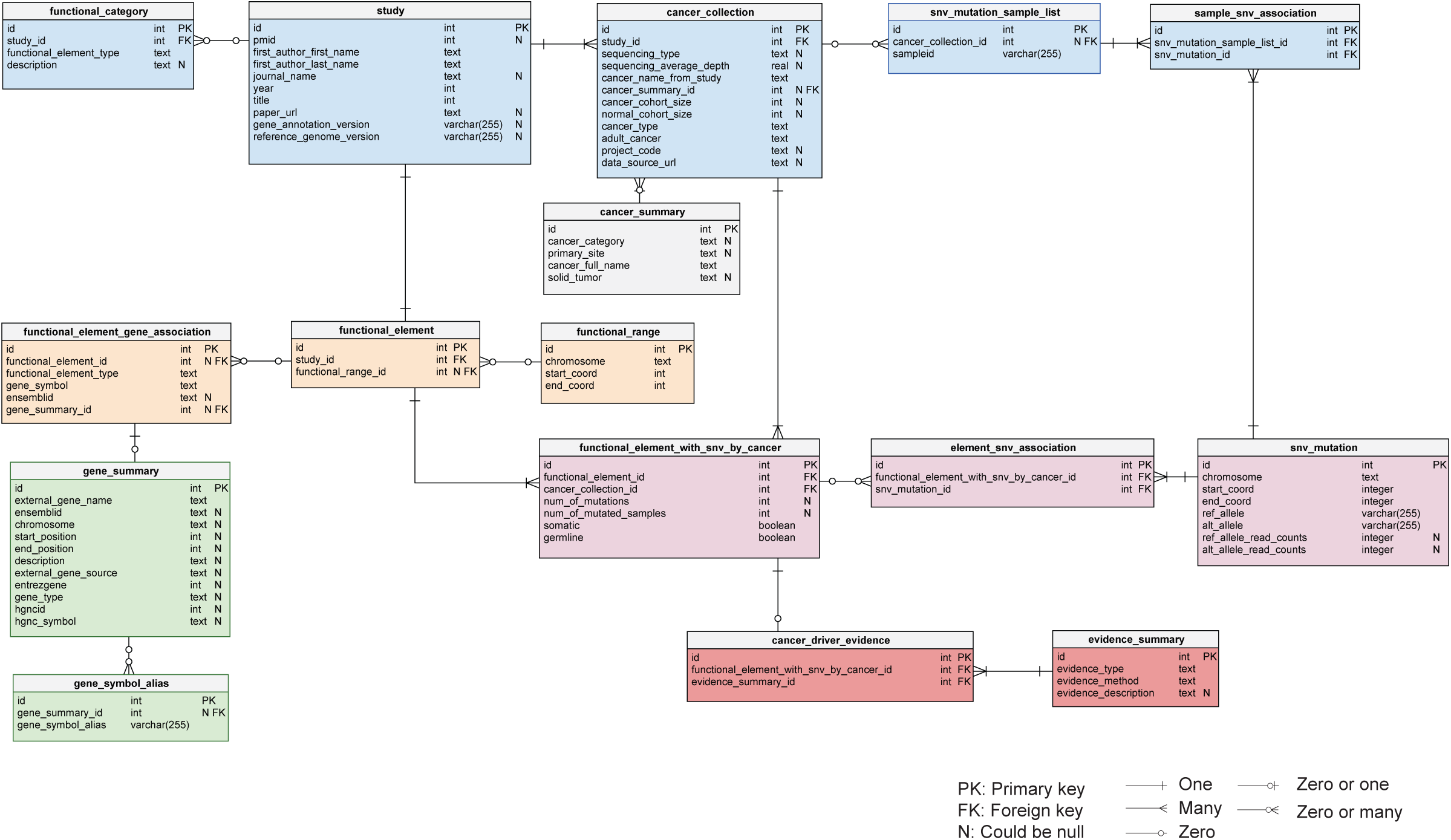
Detailed database entity-relationship diagram (ERD) in CNCDatabase.

## Notes

https://cncdatabase.med.cornell.edu/

### Competing Interest Statement

The authors have declared no competing interest.

## REFERENCES

1. Forbes, S.A., Bhamra, G., Bamford, S., Dawson, E., Kok, C., Clements, J., Menzies, A., Teague, J.W., Futreal, P.A. and Stratton, M.R. (2008) The Catalogue of Somatic Mutations in Cancer (COSMIC). In Current Protocols in Human Genetics. John Wiley & Sons, Inc., Hoboken, NJ, USA, Vol. Chapter 10, p. Unit 10.11.

2. Chakravarty, D., Gao, J., Phillips, S., Kundra, R., Zhang, H., Wang, J., Rudolph, J.E., Yaeger, R., Soumerai, T., Nissan, M.H., et al. (2017) OncoKB: A Precision Oncology Knowledge Base. JCO Precis. Oncol., 1, 1–16.

3. Griffith, M., Spies, N.C., Krysiak, K., McMichael, J.F., Coffman, A.C., Danos, A.M., Ainscough, B.J., Ramirez, C.A., Rieke, D.T., Kujan, L., et al. (2017) CIViC is a community knowledgebase for expert crowdsourcing the clinical interpretation of variants in cancer. Nat. Genet., 49, 170–174.

4. Gonzalez-Perez, A., Perez-Llamas, C., Deu-Pons, J., Tamborero, D., Schroeder, M.P., Jene-Sanz, A., Santos, A. and Lopez-Bigas, N. (2013) IntOGen-mutations identifies cancer drivers across tumor types. Nat. Methods, 10, 1081–2.

5. Huang, F.W., Hodis, E., Xu, M.J., Kryukov, G. V., Chin, L. and Garraway, L.A. (2013) Highly Recurrent TERT Promoter Mutations in Human Melanoma. Science (80-.)., 339, 957–959.

6. Bailey, S.D., Desai, K., Kron, K.J., Mazrooei, P., Sinnott-Armstrong, N.A., Treloar, A.E., Dowar, M., Thu, K.L., Cescon, D.W., Silvester, J., et al. (2016) Noncoding somatic and inherited single-nucleotide variants converge to promote ESR1 expression in breast cancer. Nat. Genet., 48, 1260–1266.

7. Gupta, R.A., Shah, N., Wang, K.C., Kim, J., Horlings, H.M., Wong, D.J., Tsai, M.-C., Hung, T., Argani, P., Rinn, J.L., et al. (2010) Long non-coding RNA HOTAIR reprograms chromatin state to promote cancer metastasis. Nature, 464, 1071–1076.

8. Liu, E.M., Martinez-Fundichely, A., Diaz, B.J., Aronson, B., Cuykendall, T., MacKay, M., Dhingra, P., Wong, E.W.P., Chi, P., Apostolou, E., et al. (2019) Identification of Cancer Drivers at CTCF Insulators in 1,962 Whole Genomes. Cell Syst., 0.

9. Cuykendall, T.N., Rubin, M.A. and Khurana, E. (2017) Non-coding genetic variation in cancer. Curr. Opin. Syst. Biol., 1, 9–15.

10. Khurana, E., Fu, Y., Colonna, V., Mu, X.J., Kang, H.M., Lappalainen, T., Sboner, A., Lochovsky, L., Chen, J., Harmanci, A., et al. (2013) Integrative Annotation of Variants from 1092 Humans: Application to Cancer Genomics. Science (80-.)., 342, 1235587.

11. Khurana, E., Fu, Y., Chakravarty, D., Demichelis, F., Rubin, M.A. and Gerstein, M. (2016) Role of non-coding sequence variants in cancer. Nat. Rev. Genet., 17, 93–108.

12. Rheinbay, E., Nielsen, M.M., Abascal, F., Wala, J.A., Shapira, O., Tiao, G., Hornshøj, H., Hess, J.M., Juul, R.I., Lin, Z., et al. (2020) Analyses of non-coding somatic drivers in 2,658 cancer whole genomes. Nature, 578, 102–111.

13. Weinhold, N., Jacobsen, A., Schultz, N., Sander, C. and Lee, W. (2014) Genome-wide analysis of noncoding regulatory mutations in cancer. Nat. Genet., 46, 1160–1165.

